# Comparison of radiography and computed tomography for condylar fracture risk assessment in Thoroughbred racehorses

**DOI:** 10.1101/2023.11.09.566089

**Authors:** S. Irandoust, L. O’Neil, C.M. Stevenson, F.M. Franseen, P.H.L. Ramzan, S.E. Powell, S.H. Brounts, S.J. Loeber, D.L. Ergun, R.C. Whitton, C.R. Henak, P. Muir

## Abstract

**Background:** Catastrophic injury has a low incidence but leads to the death of many Thoroughbred racehorses annually. Effective screening for injury risk needs to solve the false negative diagnostic sensitivity problem.

**Objectives:** To determine sensitivity, specificity, and reliability for condylar stress fracture risk assessment from fetlock digital radiographs (DR) and standing computed tomography (sCT) imaging (Asto CT Equina^®^).

**Study design:** Controlled *ex vivo* experiment.

**Methods:** A blinded set of DR and sCT images of the thoracic limb fetlock were prepared from 31 Thoroughbred racehorses and reviewed by four veterinarians. Observers evaluated the condyles and parasagittal grooves (PSG) of the third metacarpal subchondral bone (MC3) for the extent of dense bone and lucency/fissure and assigned a risk assessment grade for condylar stress fracture based on imaging findings. Sensitivity and specificity for detection of subchondral structural changes in the condyles and PSG of the third metacarpal bone and for risk assessment for condylar stress fracture were determined by comparison with a reference assessment. Agreement between each observer and the reference assessment and reliability between observers were determined. Intra-observer repeatability was also assessed.

**Results:** Intra-observer repeatability was identified for both DR and sCT imaging. Sensitivity for detection of structural change was lower than specificity for both imaging methods and all observers. For horses with a normal level of risk, observer assessment often agreed with the reference assessment. Sensitivity for risk assessment was lower than specificity for all observers. For horses with a high risk of serious injury, observers generally underestimated the level of risk. Diagnostic sensitivity of risk assessment was improved with sCT imaging, particularly for horses with elevated risk of injury. Assessment reliability was better with sCT than DR.

**Main limitations:** The *ex vivo* study design influenced DR image sets regarding limb positioning and image contrast compared with *in vivo* DR imaging.

**Conclusions:** Risk assessment through screening with diagnostic imaging is a promising approach to improve injury prevention in racing Thoroughbreds. Knowledge of sensitivity and specificity of fetlock lesion detection by DR and sCT provides critical guidance regarding development of improved screening programs for racehorses using diagnostic imaging. We found improved detection of MC3 subchondral structural change and risk assessment for condylar stress fracture with sCT *ex vivo*.

## 1 | INTRODUCTION

Risk of exercise-associated mortality and risk of catastrophic injury for Thoroughbred racehorses can be as high as 0.55% of race starts^1^ with a pooled incidence of 0.115% in flat racing.^2^ The regional incidence of third metacarpal/metatarsal (MC3/MT3) condylar stress fractures is variable but can represent 25% of catastrophic injuries in some regions such as California^3^ and are more common in the MC3 bone compared with the MT3 bone.^4^ Progressive accumulation of subchondral fatigue microdamage over time eventually leads to structural failure of the bone.^5^ This highlights the potential benefit of a screening approach to injury prevention, as misdiagnosis of the progressive fatigue damage can lead to catastrophic fractures.^6^ Due to the shape of the distal end of the MC3/MT3 bone, overlying bones, the density of the bone end, and the minimal clinical signs^7^ often associated with fatigue injury to the distal MC3/MT3, screening to identify racehorses in training with a high imminent risk of condylar stress fracture is challenging. Such early detection is important, because subsequent progression to a complete stress fracture at high speed is associated with risk of mortality, surgical intervention and serious career interruption or curtailment. Athletic performance after surgical treatment of Thoroughbreds with condylar fracture can be disappointing because of osteoarthritis, the presence of multiple condylar fractures, fracture comminution, and fractures in adjacent bones.^4,8,9^ There is substantial societal concern regarding the welfare of Thoroughbred racehorses and the problem of catastrophic musculoskeletal injury and associated risk of jockey injury is a particular focus.^10^ Screening the Thoroughbred racehorse fetlock to mitigate the risk of condylar stress fracture and the potential for serious or catastrophic injury is an area of intense interest currently.^10^

Very high cyclic loads are transferred from the proximal sesamoid bones to the condyles of the MC3/MT3 during racing.^11^ With race training, isotropic trabecular bone in the distal end of the cannon bone becomes remodeled into anisotropic bone with parallel sagittal plates extending proximally from the articular surface that are connected by thin mediolateral struts.^12–15^ Fracture is initiated through local accumulation of fatigue microdamage arising from calcified cartilage with coalescence into a macroscopic PSG fatigue crack.^5,16–18^ Once parasagittal groove (PSG) fatigue damage has compromised the mechanical integrity of the cortical shell of the distal end of the bone, a condylar stress fracture may propagate proximally through the thin medial-lateral struts in the trabecular bone,^16^ although initial propagation may be delayed by focally increased adaptive sclerosis of the distal MC3/MT3 in response to high strain cyclic loading. An associated remodeling response is typical and may be seen as localized bone lysis in the PSG of the subchondral bone radiographically.^6,16–19^ Subchondral fatigue cracks may also propagate obliquely to form a palmar/plantar osteochondral disease (POD) lesion which occurs primarily within the condyle. The advanced POD lesion is typically a saucer shaped subchondral bone fatigue injury (SBI) that is also initiated by accumulation of articular microcracks, initially in the calcified cartilage zone. Although uncommon, condylar fractures that arise more abaxially in the joint surface can be initiated by subchondral fatigue injury associated with a POD lesion.^17,20,21^ POD lesions are not typically evident with DR unless pathologic changes are severe; their role in risk of catastrophic injury is currently considered to be minimal but is not fully determined.^22,23^ Incipient condylar fracture can be diagnosed by planar DR through identification of altered radiodensity in the PSG region.^6^ However, the reliability of digital radiography (DR) for detection of concerning SBI in the PSG that precipitates condylar stress fracture is unclear. Focal PSG SBI in some horses may not be detectable by DR,^24^ and the consequence of a false negative diagnosis may be catastrophic.^6^ Standing computed tomography (sCT) may enhance identification of fatigue-induced structural change.^24,25^ However, the usefulness of both modalities is contingent upon equipment, technique, and user interpretation. Currently, there is little published information available to guide clinicians in the application of DR or sCT for identification of Thoroughbred racehorses with concerning PSG SBI associated with increased risk of imminent catastrophic injury. The observation that PSG SBI lesions are much more common in the contralateral limbs of Thoroughbred racehorses with catastrophic condylar stress fracture compared with horses that are not catastrophically injured^26,27^ highlights the need to address this gap in knowledge. In a recent study, 5 of 13 horses with condylar stress fractures had small focal PSG SBI lesions in the contralateral limb, whereas this finding was not observed in any of the 8 control horses.^26^

The flexed dorsopalmar or plantarodorsal radiographic project is critical for identification of this type of radiographic change using DR.^6^ Optimal radiographic technique is important for diagnosis of incipient condylar fractures using DR and current best practice uses multiple flexed projections.^28^ The growing application of cross-sectional imaging methods to the Thoroughbred fetlock over time has encouraged refinement of DR technique and interpretation in recent years^29^ and promises improved screening of Thoroughbred racehorses.^10^

To better understand the role of DR and sCT in detection of incipient condylar fracture, we compared detection of SBI lesions in the Thoroughbred racehorse fetlock *ex vivo* using the two imaging methods. We hypothesized that sensitivity and specificity of detection of concerning PSG SBI would be influenced by imaging method and by the clinical experience of the observer.

## 2 | MATERIALS AND METHODS

Because condylar stress fractures are most common in the thoracic limb, *ex vivo* imaging in this study was limited to the thoracic limb. This helped generate a standardized image set for blinded evaluation by a panel of observers against a gold standard reference for determination of sensitivity and specificity of structural feature detection and clinical risk assessment for condylar stress fracture.

### 2.1 | Sample population of Thoroughbred racehorses

Entire distal limb specimens were obtained from 31 Thoroughbred racehorses that died or were euthanatized for reasons unrelated to the present study during athletic training and racing and were given for use in this work. Horses were euthanatized humanely by a veterinarian at the racetrack because of catastrophic injury. Limbs were transected at the level of the carpus, sealed in plastic bags, and stored at −20C until needed. Limbs were thawed to room temperature before use.

### 2.2 | Fetlock digital radiography

DR imaging used a VetRocket Xray generator (HF100/30+ generator, Min-Xray Inc., Northbrook, IL) and panel (CDXI-31, Canon Electron Tubes and Devices Co., Ltd. and Sumito Corporation of the Americas, Rosemont, IL). For each study fetlock, the following carefully collimated radiographic projections were made using a standard protocol^28^ and a positioning jig: 1) Flexed latero-medial; 2) Flexed dorsal-palmar with five projections including horizontal and ±7.5 and ±15 degrees. Radiographic exposure settings were set at 64kVp and 4.25mAs and a focal distance of 91.4cm (36 inches). Positioning in each image was checked for technical errors^21,28^ before the DICOM images were saved to a network drive. Each image set was anonymized for evaluation and a screenshot flexed dorsopalmar image of each limb was also prepared to indicate the lateral aspect of the fetlock.

Blinded DR image sets were evaluated for the extent of dense subchondral bone in the lateral and medial condyles and PSGs (0 - absent, 1 - mild [<33% of the subregion], 2 - moderate [33-66% of the subregion], 3 - severe [>66% of the subregion]). Images were also evaluated for the presence of a subchondral lucency/fissure in the lateral and medial condyles and PSGs (0 - absent, 1 - poorly defined, no surrounding increased density, 2 - poorly defined, with surrounding increased density, 3 - well defined, no surrounding increased density, 4 - well defined, with surrounding increased density) (**Supplementary File S1)**. Each horse was also assigned a risk assessment grade based on structural features from diagnostic imaging (**Table 1**, **Supplementary File S1**). For the purposes of risk assessment screening using diagnostic imaging, ‘overt stress fracture’ was defined as a subchondral structural abnormality associated with sufficient fatigue damage where the observer considered the mechanical properties of the bone compromised based on current knowledge in the field.^6,16,19,24,26,30,31^

**TABLE 1.** Overall assessment of immediate risk of condylar stress fracture in the racing Thoroughbred.

| **Risk Category** | **Clinical Recommendation** |
| --- | --- |
| Normal | Standard risk. Continue training and racing |
| Indeterminate | Further imaging recommended |
| Strong likelihood of stress crack from subchondral fatigue injury | Discontinue fast training and further imaging is recommended |
| Stress crack present from subchondral fatigue injury | Discontinue training and further imaging recommended to assess significance |
| Overt stress fracture from fatigue injury | Rehabilitation or surgery is required |

### 2.3 | Fetlock standing computed tomography

Each limb specimen was scanned frozen to enable specimen preservation and to minimize tissue exposure to freeze-thaw cycles. Scanning frozen does not alter CT imaging of osseous structures. sCT imaging was performed at high-resolution using 0.55mm contiguous slices using a robotic CT scanning system which has the capability of performing vertical scanning of limb pairs as well as horizontal scanning of the head/neck in the standing sedated horse. The 24-slice helical Equina^®^ scanner (Asto CT Equina^®^) is a fixed installation and has a 240V single phase 30A uninterruptable power supply. Scanning was performed using an exposure of 160kVp and 8mA at 1 second per 360° revolution using 24 detector rows with a variable helical pitch, typically 0.55. Slice acquisition rate is 36 slices/sec with an image acquisition matrix of 1024×1024 and a resolution at isocenter of 0.75mm. Maximum scanning distance is 100cm vertically at 2cm/second.^25^ No scout images are needed. Scanning was performed in a horizontal direction with the limbs placed on the positioning table normally used for head scanning in the standing horse. DICOM images were then stored on a network drive. Each image set was anonymized for evaluation using the Horos DICOM viewer and a screenshot transverse image at the level of the MC3 condyles was also prepared to indicate the lateral aspect of the fetlock.

Blinded sCT image sets were evaluated by multiplanar reconstruction for increased subchondral bone density in the lateral and medial condyles and PSGs using a 45° oblique dorsoproximal palmarodistal reconstruction perpendicular to the palmar joint surface. The sagittal plane was positioned over the region of interest for grading using an established scale.^32^ Although microdamage cannot be detected in living horses with current imaging methods, the extent of radiographic subchondral bone sclerosis is correlated with microdamage.^33,34^ Also, the thickness of the subchondral plate may be linked to risk of condylar stress fracture.^26^ In each PSG, the subchondral plate thickness (mm) was measured perpendicular to the articular surface of the condyles in the same 45° degree oblique dorsal proximal to palmar distal reconstruction as a measure of subchondral sclerosis and associated fatigue microdamage (**Supplementary File S1**).^26^ Images were also evaluated for the presence of a subchondral lucency/fissure in the lateral and medial condyles and PSGs (0 - absent, 1 poorly defined, no surrounding increased density, 2 - poorly defined, with surrounding increased density, 3 - well defined, no surrounding increased density, 4 - well defined, with surrounding increased density) as for the DR images (**Supplementary File S1**). Each horse was also assigned a risk assessment grade based on structural features from diagnostic imaging (**Table 1, Supplementary File S1**). Parasagittal area (mm^2^) of the subchondral lucency was measured when identified.^30^

### 2.4 | Visual examination of the distal articular surface of the third metacarpal bone and photography

Pathological examination used an established approach.^33^ After diagnostic imaging, all non-cartilaginous soft tissue was removed from each MC3 bone. The shaft was cut with a band saw in the mid-diaphysis, 16.5cm (6.5 inches) proximal to the dorsal articular margin of the distal end of the bone. The second and fourth metacarpal bones were removed. The articular surface at the distal end of the bone was photographed and the articular cartilage and any remaining soft tissues were then macerated using 0.15–0.2 M NaOH at 37°C. The solution was changed as needed every 1 to 3 days. Once the soft tissues were removed, additional photographs of the subchondral bone of the distal articular surface were made.

Severity of fatigue injury to the distal MC3 articular surface in the lateral and medial condyles and PSGs was graded before and after removal of hyaline articular cartilage by maceration (0 - absent, 1 - mild, 2 - moderate, 3 - severe). Lesions typically have a linear shape in the PSG and a circular shape in the condyle. Features assessed included discoloration of the cartilage or subchondral bone of the palmar joint surface, ulceration/collapse of the cartilage, or presence of subchondral defect.^35–37^

### 2.5 | Radiographic and pathologic image evaluation

Four blinded observers were used for image assessment with differing clinical experience. A boarded radiologist with extensive experience working with Thoroughbred racehorses (SEP, Observer 1), a boarded large animal surgeon (SHB, Observer 2), and a second boarded radiologist with equine emphasis (SJL, Observer 3) evaluated the DR and sCT image sets. A primary care veterinarian with extensive experience working with Thoroughbred racehorses in training (PHLR, Observer 4) also evaluated the DR image set. In the randomized blinded DR and sCT image sets, images from one Thoroughbred were included five times to determine intra-observer variation.

Review of sCT combined with visual assessment of the joint surface of the distal end of the MC3 bone to further assess fatigue damage to the joint surface was considered the gold standard reference interpretation,^20^ because observation of the joint surface provides direct detailed assessment for the presence of PSG subchondral fatigue cracks. This assessment was undertaken by an experienced observer (PM) who evaluated the blinded sCT image sets and generated assessment data for structural change and risk assessment using the study assessment form (**Supplementary File S1**). Reference assessment of subchondral sclerosis was based on sCT evaluation. Subsequently, joint surface photographs were evaluated from each horse to generate assessment scores for cartilage and subchondral bone injury. Finally, sCT images were reevaluated together with the joint surface photographs to develop an updated gold standard risk assessment score for imminent injury for each horse and confirm sCT assessments of SBI.

### 2.5 | Data analysis

The image sets that were repeatedly scored 5 times were condensed to a single value for each observer and each modality for inter-observer comparisons. The median value for the categorical scores, the mean value for continuous measurements, and the majority risk assessment grade were used as the final entry for the repeated horse in the data set for each observer in each modality. Contingency tables were constructed using Prism v10.0 (GraphPad Software, Boston, MA) for each imaging method and for each clinician observer to determine sensitivity and specificity for detection of structural changes to distal MC3 subchondral bone and condylar stress fracture risk assessment against the reference assessment. The Fisher’s exact test was used to determine significant differences between feature identification from each observer and the reference interpretation for DR imaging, sCT imaging, and the associated risk predictions for each horse. Results were considered significant at *P*<0.05.

The intraclass correlation coefficient (ICC) (*r*) statistic was calculated^38^ using MATLAB R2023a (MathWorks, Natick, MA) to estimate intra-observer repeatability, agreement between each observer and the reference assessment and inter-observer reliability in rating structural features and condylar stress fracture risk assessment. For interpretation of agreement between each observer and the reference assessment and inter-observer reliability, <0.5 was considered poor reliability, ≥0.5 to <0.75 was considered moderate reliability, ≥0.75 to <0.9 was considered good reliability, and ≥0.9 was considered excellent reliability.

## 3 | RESULTS

### 3.1 | Postmortem findings

The horses of the study had a range of joint pathologic change from mild to severe (**Supplementary Figures S1-2**). After initial evaluations made using sCT, risk assessments for 8 horses were revised after examination of the articular surface photographs before and after cartilage removal. Risk assessment for these 8 horses was updated from *Normal* based on sCT alone to *Stress crack present from subchondral bone injury* after postmortem evaluation identified small macroscopic cracks in the PSG (**Figure 1**). Of these 8 horses, 5 were diagnosed as *Normal* by all 4 observers. Assessment of risk was variable for the other 3 horses across the observers.

**FIGURE 1.**
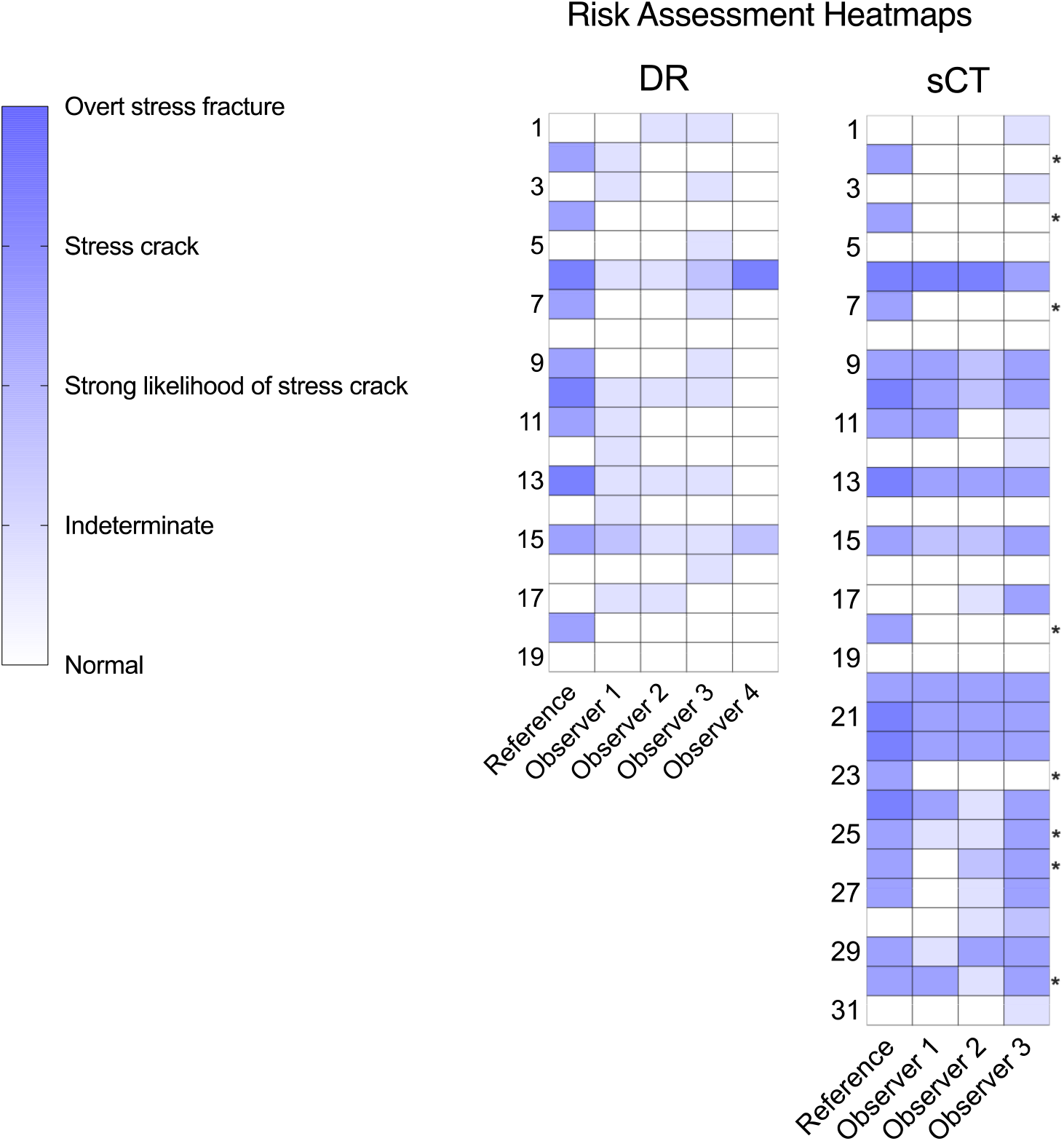
Condylar stress fracture risk assessment from digital radiography (DR) and standing CT (sCT) from a population of racing Thoroughbreds. Reference – Gold standard reference assessment using sCT combined with examination of the joint surface before and after digestion of articular cartilage. For many horses, risk was assessed at a lower level than the reference assessment, particularly for DR. Most horses with elevated fracture risk were identified by multiple sCT observers. * Risk assessment for these 8 horses was updated from *Normal* based on sCT alone to *Stress crack present from subchondral bone injury* after postmortem evaluation.

### 3.2 | Intra-observer repeatability

Intra-observer repeatability is reported in **Tables 2–4**. Variation in assessment of structural change was found for all observers with DR (**Table 2**). All readings were within one grade, except for one observer with one density grade where the difference was two grades (**Table 2**). Lucency/fissure in the PSG with DR imaging was consistently reported as absent (0) by Observer 3. Variation was found in the continuous variables subchondral plate thickness and PSG lucency parasagittal area measured from sCT by all observers as well as the categorical variables (**Table 3**). The magnitude of intra-observer variation for subchondral plate thickness ranged from 6-23% across observers and 17-157% for PSG lucency parasagittal area.

**TABLE 2.** Intra-observer repeatability in detection of structural change reported by digital radiography for the racing Thoroughbred with five blinded repeats of the image set.

| Observer | Extent of dense metacarpal subchondral bone |  |  |  | Focal lucency/fissure |  |  |  |
| --- | --- | --- | --- | --- | --- | --- | --- | --- |
|  | LC | LPSG | MPSG | MC | LC | LPSG | MPSG | MC |
| 1 | 1 (1, 2) | 1 (0, 1) | 0 (0, 1) | 2 (1, 2) | 0 (0) | 1 (1, 2) | 1 (1) | 0 (0) |
| 2 | 1 (0, 2) | 1 (1, 2) | 2 (1, 2) | 1 (1, 2) | 0 (0) | 0 (0, 1) | 1 (1, 2) | 0 (0) |
| 3 | 1 (1) | 1 (0, 1) | 0 (0, 1) | 1 (1) | 0 (0) | 0 (0) | 0 (0) | 0 (0, 1) |

**Note.** Data are reported as median (range). Observer 4 reported poor image quality for all 5 of the DR image sets and did not score any of them. The extent of dense subchondral bone was evaluated as 0 – absent, 1 – mild, 2 – moderate, 3 – severe and focal lucency/fissure was evaluated as 0 – absent, 1 – poorly defined, no surrounding increased density, 2 – poorly defined with surrounding increased density, 3 – well defined, no surrounding increased density, 4 – well defined with surrounding increased density (**Supplementary Figure S1**). LC – lateral condyle, LPSG – lateral parasagittal groove, MPSG – medial parasagittal groove, MC – medial condyle.

**TABLE 3.** Intra-observer repeatability in detection of structural change reported by standing computed tomography for the racing Thoroughbred with five blinded repeats of the image set.

| Observer | Extent of dense metacarpal subchondral bone |  |  |  | Subchondral plate thickness (mm) |  | Focal lucency/fissure |  |  |  | Focal lucency parasagittal area (mm<sup>2</sup>) |  |
| --- | --- | --- | --- | --- | --- | --- | --- | --- | --- | --- | --- | --- |
|  | LC | LPSG | MPSG | MC | LPSG | MPSG | LC | LPSG | MPSG | MC | LPSG | MPSG |
| Reference | 1 (1, 2) | 2 (1, 2) | 2 (2) | 1 (1, 2) | 17±2.35 | 19±1.30 | 0 (0) | 4 (4) | 3 (3) | 0 (0) | 24 (21, 28) | 56 (52, 61) |
| 1 | 2 (2, 3) | 2 (1, 2) | 2 (1, 2) | 2 (2, 3) | 9±1.10 | 9±1.14 | 0 (0, 3) | 4 (3, 4) | 4 (0, 4) | 0 (0, 3) | 15 (7, 18) | 26 (24, 28) |
| 2 | 2 (2, 3) | 2 (2, 3) | 2 (2, 3) | 2 (2, 3) | 21±1.29 | 21±2.54 | 0 (0) | 4 (4) | 3 (1, 4) | 0 (0) | 38 (28, 44) | 27 (19, 36) |
| 3 | 2 (2) | 2 (2, 3) | 2 (2, 3) | 2 (2) | 21±3.27 | 21±4.86 | 0 (0) | 4 (4) | 4 (4) | 0 (0) | 13 (11, 14) | 14 (11, 18) |

**Note.** Data are reported as median (range) or mean±standard deviation. The extent of dense subchondral bone was evaluated as 0 - absent, 1 - mild, 2 - moderate, 3 - severe and focal lucency/fissure was evaluated as 0 - absent, 1 - poorly defined, no surrounding increased density, 2 - poorly defined with surrounding increased density, 3 – well defined, no surrounding increased density, 4 - well defined with surrounding increased density (**Supplementary Figure S1**). LC - lateral condyle, LPSG – lateral parasagittal groove, MPSG - medial parasagittal groove, MC - medial condyle. Observer 4 did not evaluate sCT image sets.

For condylar stress fracture risk assessment, the reference assessment of *Overt stress fracture from fatigue injury* using sCT and joint surface examination was the same for all 5 repeats. For DR, two observers also consistently graded the repeats, whilst the assessment of the third observer varied across two levels of risk (**Table 4**, **Figure 2**). For sCT, one observer consistently graded the repeats, whilst the assessments of the other two observers varied across two or three levels of risk (**Table 4**).

**TABLE 4.** Intra-observer repeatability in condylar stress fracture risk assessment determined by digital radiography (DR) and standing computed tomography (sCT) for the racing Thoroughbred with five blinded repeats of the image set.

| Risk Category | Reference | Observer 1 |  | Observer 2 |  | Observer 3 |  |
| --- | --- | --- | --- | --- | --- | --- | --- |
|  |  | DR | sCT | DR | sCT | DR | sCT |
| Normal |  |  |  |  |  |  |  |
| Indeterminate |  | 3 | 1 | 5 |  | 5 |  |
| Strong likelihood of stress crack from subchondral fatigue injury |  | 2 |  |  | 1 |  |  |
| Stress crack present from subchondral fatigue injury |  |  | 4 |  | 4 |  | 5 |
| Overt stress fracture from fatigue injury | 5 |  |  |  |  |  |  |

**Note.** Observer 4 reported poor image quality for all 5 of the DR image sets and did not score any of them. Clinical recommendations for risk assessment are reported in **Table 1**. The reference assessment was derived from standing computed tomography images combined with pathological assessment of the articular surface of the distal end of the third metacarpal bone.

**FIGURE 2.**
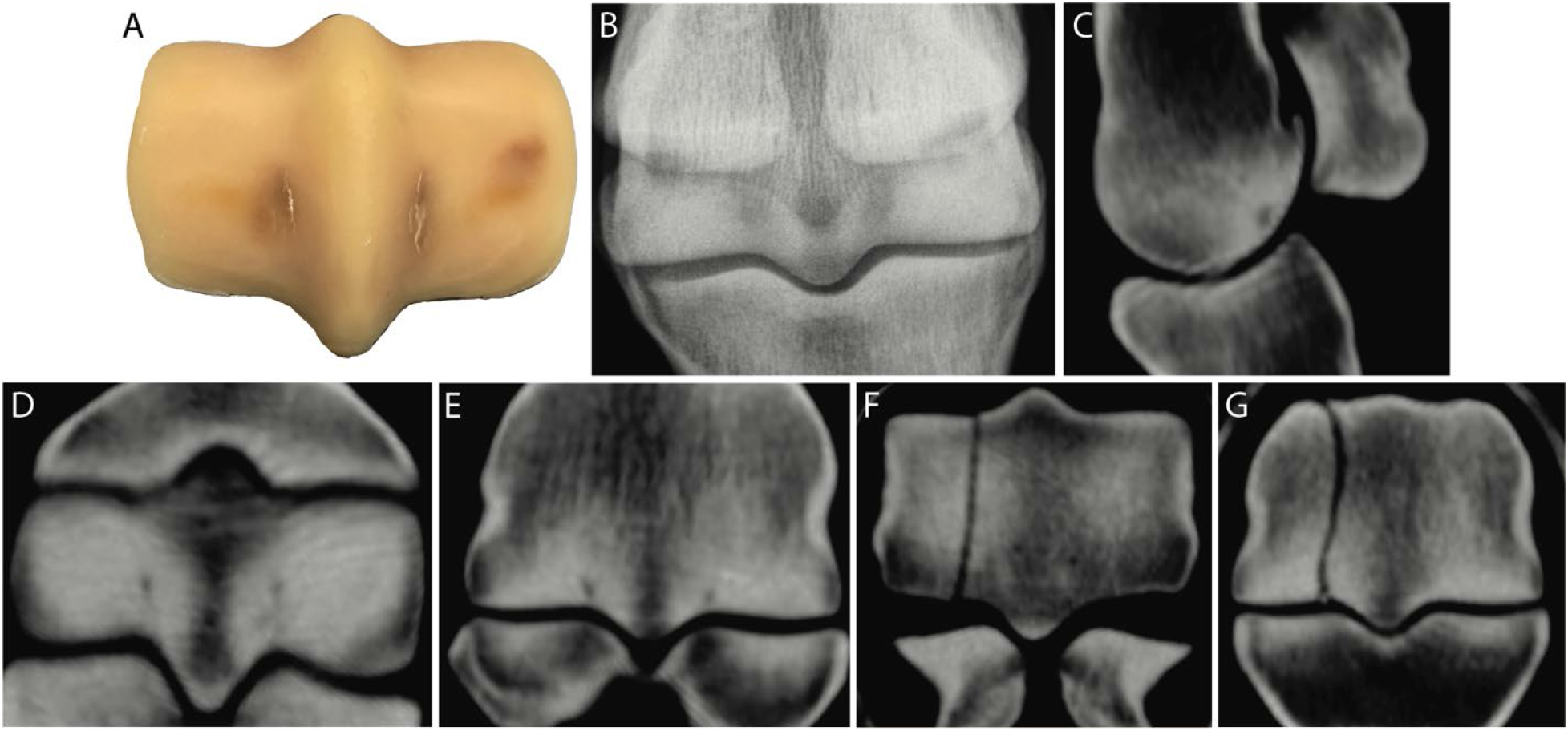
Postmortem (**A**), digital radiography (**B**), and standing computing tomography (sCT) images (**C-G**) from a racing Thoroughbred with biaxial parasagittal groove (PSG) focal subchondral bone lucencies because of fatigue injury (Horse #22, Figure 1). The lucencies seen on sagittal (**C**), transverse (**D**), and dorsal (**E**) sCT reconstructions correspond to the linear PSG subchondral fatigue cracks seen in A. Contralateral condylar stress fracture seen in transverse (**F**) and dorsal (**G**) reconstructions supports the reference assessment of *Overt stress fracture from fatigue injury*.^24,26^ **Note**. Lateral or dorsal to the left.

### 3.3 | Detection of MC3 subchondral bone injury using DR

Observer 1 did not assess image sets from 3 horses because image quality was felt to affect diagnostic interpretation. Similarly, Observer 4 did not assess image sets from 11 horses including the horse used for the repeated image set to evaluate intra-observer variation. Observer 2 and 3 evaluated images for all 31 horses. Consequently, for estimation of sensitivity, specificity, and intraclass correlation, image sets from 19 of 31 horses were used for detailed analysis.

Significant differences from the reference assessment of the extent of dense subchondral bone across the observers was most evident for bones with absent or mild changes (**Supplementary File 3**, **Figure 3**). Specificity was above 65% for detection of PSG lucency and 100% for the detection of condylar lucency for all observers (**Figure 4**). Sensitivity for detection of structural changes in subchondral bone was variable between observers (**Figure 4**). Observers 3 and 4 had low sensitivity for detection of PSG lucency/fissure.

**FIGURE 3.**
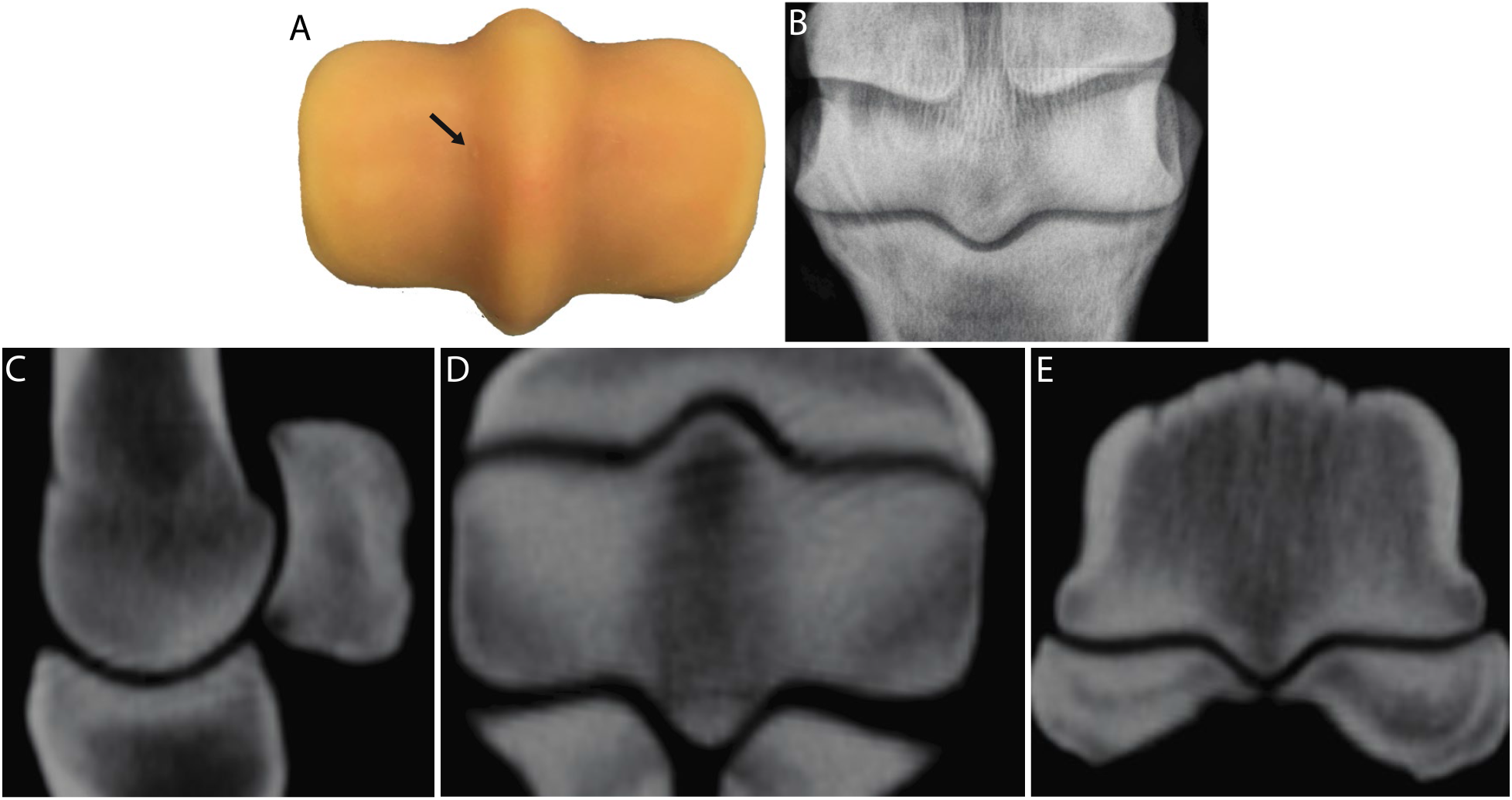
Postmortem (**A**), digital radiography (**B**), and standing computing tomography (sCT) images (**C-E**) from a racing Thoroughbred with mild subchondral injury (Horse #23, Figure 1). A very small subchondral defect is evident in the lateral PSG that was not detectable by DR or sCT. Little modeling of the subchondral plate is evident radiographically. The reference assessment was *Stress crack* while observers interpreting sCT imaging provided an assessment of *Normal*. **Note**. Lateral or dorsal to the left.

**FIGURE 4.**
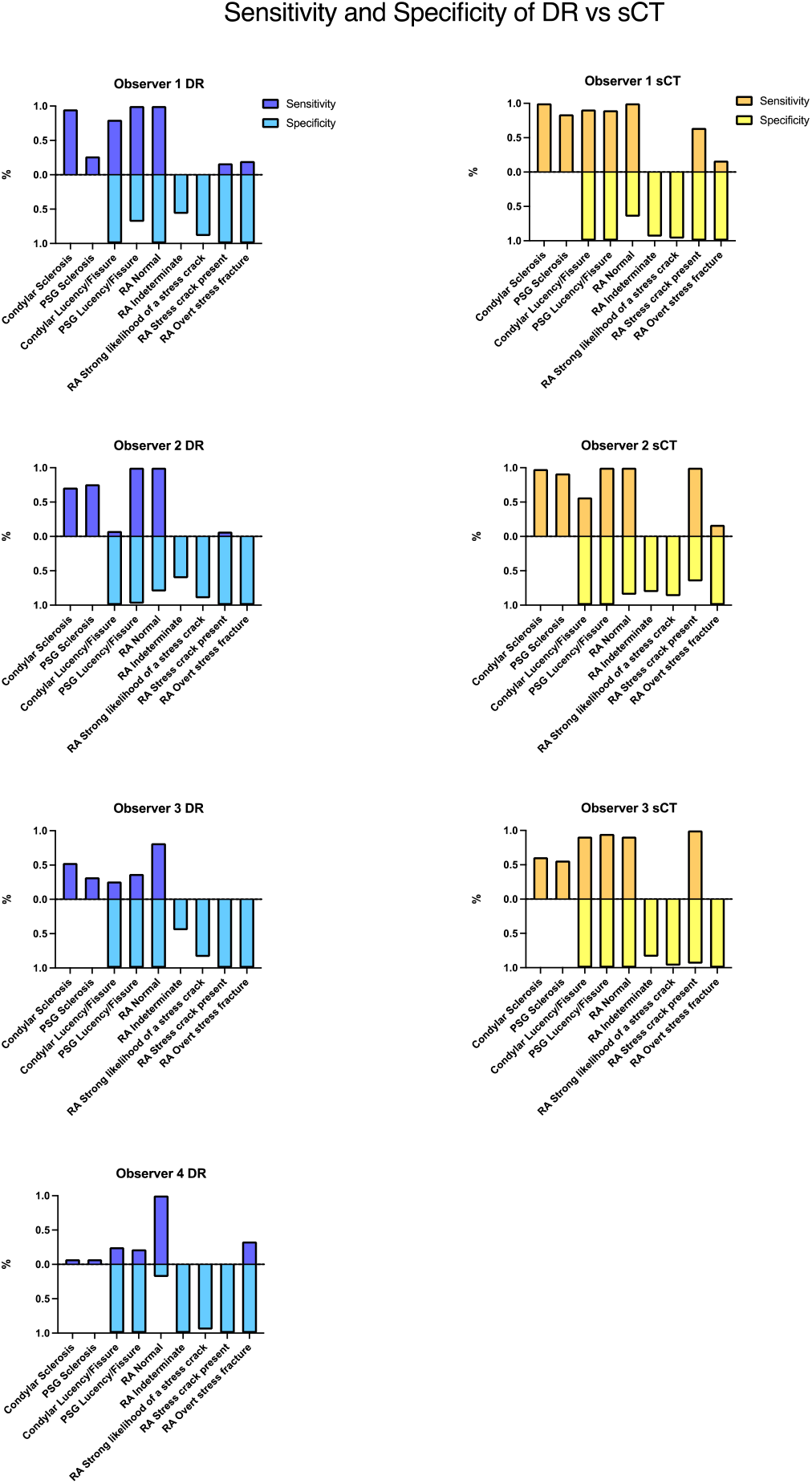
Sensitivity and specificity for detection of joint fatigue injury and condylar stress fracture risk assessment using digital radiography (DR) and standing computed tomography (sCT). Sensitivity was not reported for the *Indeterminate* and *Strong likelihood of a stress crack* categories as the reference assessment did not record any horses in these categories. Specificity could not be estimated for detection of condylar or parasagittal groove (PSG) sclerosis as the gold standard reference assessment was always yes for the detection of these changes. RA - risk assessment.

### 3.4 | Detection of MC3 subchondral bone injury using sCT

The three observers that evaluated sCT images assessed image sets from all 31 horses. Twenty one percent of the observations (10/48) differed significantly from the reference assessment of the extent of dense subchondral bone across the observers were identified (**Supplementary File 3**). Significant differences were most often identified for assessment of the extent of dense PSG subchondral bone with absent or mild change (**Supplementary File 3**). Specificity for detection of PSG and condylar lucency was 100%. Sensitivity for detection of PSG or condylar lucency was lower than specificity across the observers, except for detection of PSG lucency by Observer 2 (**Figure 4**).

Observer 1 did not measure the PSG subchondral plate thickness in one horse. Some bias from the reference measurements were identified, particularly for Observers 1 and 3 (**Supplementary Figure S3**). Both positive and negative bias amongst the observers was identified.

For measurement of sagittal plane PSG subchondral lucency area, data points were excluded if they spanned the PSG and the condyle or were present only in the condyle (**Figure 5**). All observers demonstrated close agreement with the reference assessment for the presence of a lucency (**Supplementary Table S1**). Some bias from the reference measurements was also observed for measurement of parasagittal groove subchondral lucency area in the sagittal plane. Measurements by Observers 1 and 3 showed positive bias from the reference measurements (**Supplementary Figure S4**). Measurement bias was also more variable than estimates of PSG subchondral plate thickness.

**FIGURE 5.**
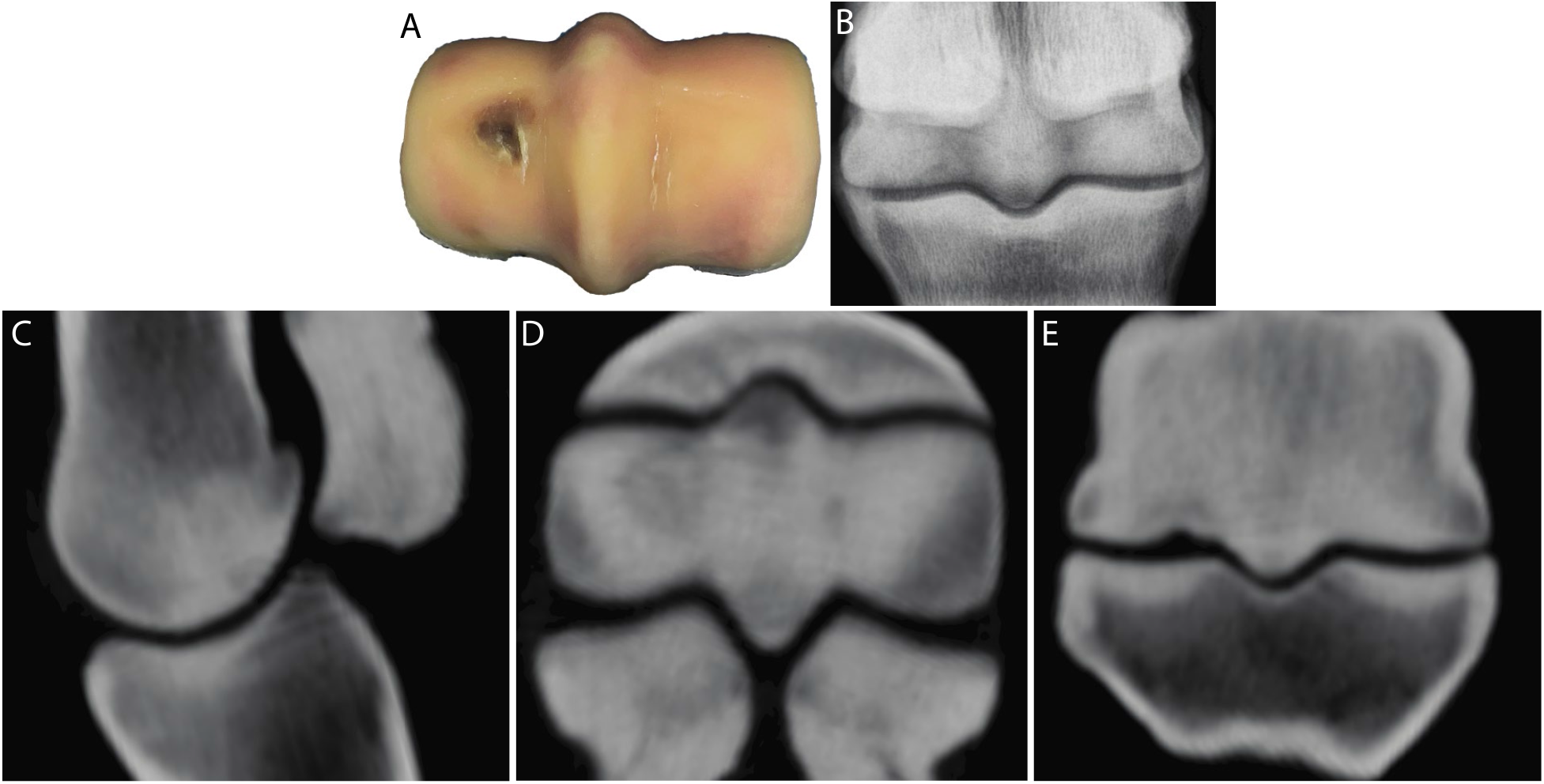
Postmortem (**A**), digital radiography (DR) (**B**), and standing computing tomography (sCT) images (**C-E**) from a racing Thoroughbred with severe subchondral changes (Horse #6, Figure 1). A large lateral palmar osteochondral disease (POD) lesion spanning the condyle and PSG together with a medial PSG subchondral lucency is evident on the sagittal (**C**), transverse (**D**) and dorsal (**E**) sCT. This fatigue damage to the joint surface corresponds with the changes evident on visual inspection (**A**). Subchondral structural changes are also evident with flexed dorsopalmar radiography (**B**) but more difficult to identify. The reference assessment was *Overt stress fracture from fatigue injury.* A lower risk assessment rating was provided by 3 of 4 observers interpreting DR imaging. Concordance with the reference assessment was improved with sCT imaging. **Note**. Lateral or dorsal to the left.

### 3.5 | Reliability of detection of MC3 subchondral bone injury using DR and sCT

For agreement with the reference measurements, the range of *r* for DR assessment was 0.0011 to 0.62 across all categories. Correlation coefficients were generally below 0.5 (**Figure 6A**). With DR, Observer 1 achieved the highest overall agreement with the reference assessment and Observer 3 achieved the lowest overall (**Figure 6A**). The range of *r* for sCT was 0.49 to 0.82 (**Figure 6A**). Correlation coefficients were generally between 0.50 and 0.75. With sCT, Observer 1 was the most consistent across the assessment variables. For agreement between observers the range of *r* for DR was 0.29 to 0.48 and for sCT was 0.65 to 0.72 (**Figure 6B**).

**FIGURE 6.**
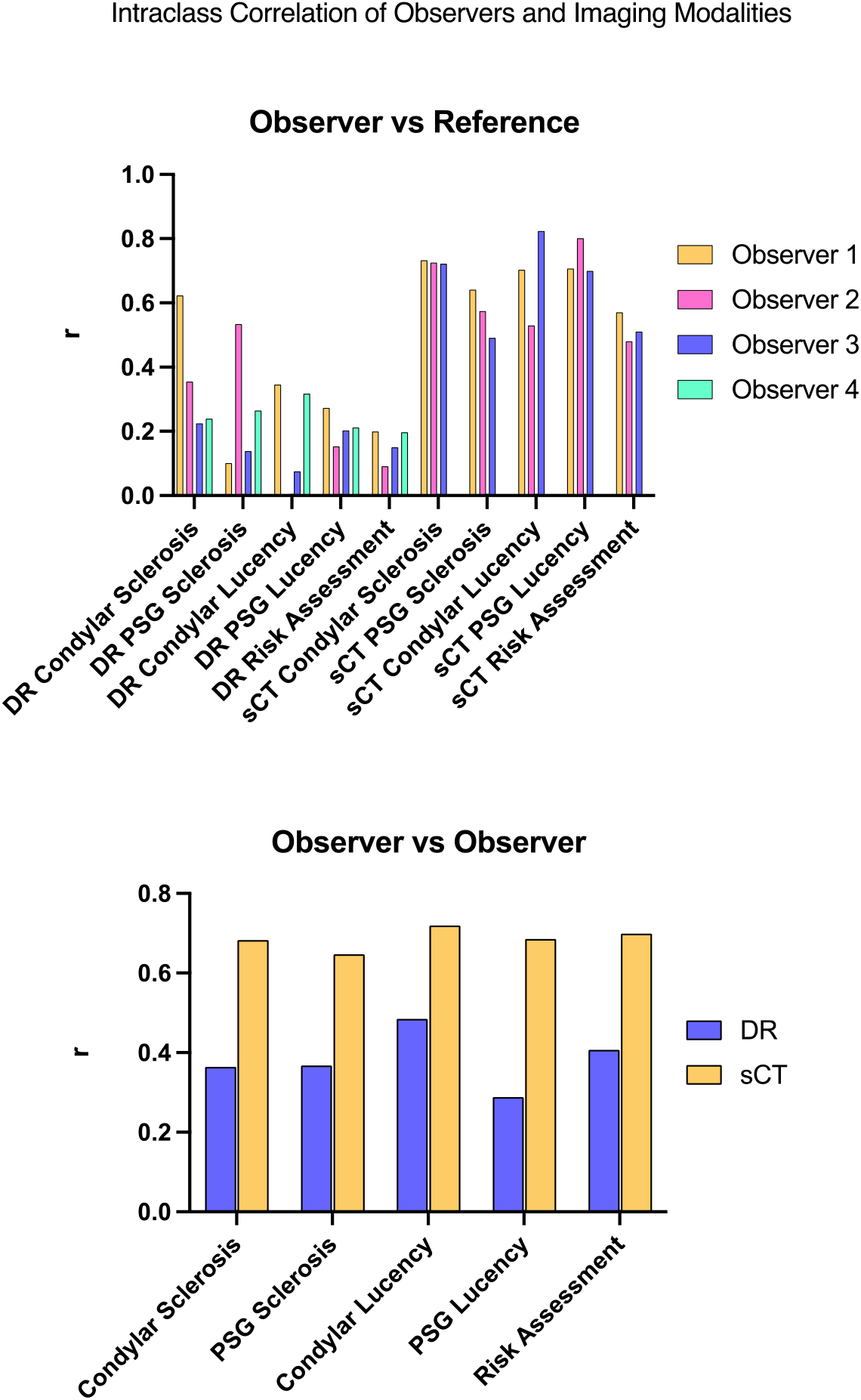
Agreement between diagnostic imaging observers estimated by the intraclass correlation coefficient statistic yield higher correlation coefficients for standing computed tomography (sCT) compared with digital radiography (DR) for both detection of structural change in the subchondral bone and for condylar stress fracture risk assessment. (**A**) Agreement between observer estimation with the gold standard reference. (**B**) Agreement between observer estimation.

### 3.6 | Accuracy of risk assessment for PSG subchondral fatigue injury with DR and sCT

For the horse used for assessment of intra-observer repeatability, the category of *Indeterminate* was most often assigned from the DR image sets (**Table 4**, **Figure 2**) different from the reference assessment of *Overt stress fracture from fatigue injury*. In contrast, a risk assessment of *Stress crack present from subchondral fatigue injury* was most often assigned from sCT for this horse (**Table 4**).

Sensitivity was not reported for the *Indeterminate* and *Strong likelihood of a stress crack* categories as the reference assessment did not record any horses in these categories (**Tables 5–6**). For horses with a standard level of risk rated as *Normal*, observer risk assessment often agreed with the reference assessment. Specificity for risk assessment was much higher than sensitivity for all observers (**Figure 4**). The only category with 100% sensitivity with DR was *Normal* for 2 observers. For horses with elevated risk assessed to have a *stress crack present from subchondral fatigue injury* or an *overt stress fracture from fatigue injury* observers generally underestimated the level of risk. Horses with elevated risk were most accurately assessed by Observer 4 with associated higher diagnostic sensitivity (**Table 5**, **Figure 4**). For each observer, diagnostic sensitivity of risk assessment was improved with sCT imaging, particularly for horses with elevated risk of injury.

**TABLE 5.** Accuracy of condylar stress fracture risk assessment in the racing Thoroughbred using digital radiography.

| **Risk Category** | **Reference** | **Observer**<br>**1** | **Observer**<br>**2** | **Observer**<br>**3** | **Observer**<br>**4** |
| --- | --- | --- | --- | --- | --- |
| Normal | 9 | 9 | 13 | 9 | 17 |
| Indeterminate |  | 9 | 6 | 9 |  |
| Strong likelihood of stress crack from subchondral fatigue injury |  | 1 |  | 1 | 1 |
| Stress crack present from subchondral fatigue injury | 7 |  |  |  |  |
| Overt stress fracture from fatigue injury | 3 |  |  |  | 1 |

**Note.** Analysis was based on the 19 horses that all 4 observers assessed. The reference assessment was based on evaluation of standing computed tomography images combined with pathological assessment of the articular surface of the distal end of the third metacarpal bone.

**TABLE 6.** Accuracy of condylar stress fracture risk assessment in the racing Thoroughbred using standing computed tomography.

| **Risk Category** | **Reference** | **Observer**<br>**1** | **Observer**<br>**2** | **Observer**<br>**3** |
| --- | --- | --- | --- | --- |
| Normal | 11 | 18 | 15 | 10 |
| Indeterminate |  | 2 | 6 | 5 |
| Strong likelihood of stress crack from subchondral fatigue injury |  | 1 | 4 | 1 |
| Stress crack present from subchondral fatigue injury | 14 | 9 | 5 | 15 |
| Overt stress fracture from fatigue injury | 6 | 1 | 1 |  |

**Note.** Analysis was based on all 31 horses that all 3 observers assessed. The reference assessment was based on evaluation of standing computed tomography images combined with pathological assessment of the articular surface of the distal end of the third metacarpal bone.

Regarding agreement with the reference analysis, the range of *r* was 0.09 to 0.20 for DR assessment and 0.48 to 0.57 for sCT (**Figure 6**). For agreement between readers, *r*=0.41 for DR and *r*=0.70 for sCT (**Figure 6**, **Supplementary Figure S5**).

## 4 | DISCUSSION

At the population level, achieving substantial reduction in incidence of catastrophic injury continues to be a challenging problem to address, in part because gait changes associated with fetlock PSG SBI may only be evident at full racing speed and pre-race veterinary inspection may not reveal concerning clinical signs.^7,39^ Interactions between limb loading from athletic activity and racing surface properties, microdamage accumulation and bone remodeling appear complex. Consequently, there is increasing focus on screening of the individual horse using diagnostic imaging to identify at-risk horses and more research is needed.^10^ Any effective screening test to prevent an acute potentially catastrophic event should be highly sensitive (few false negatives) for detection of at-risk horses. Test specificity (few false positives) is also impactful, because the Thoroughbred racehorse industry will not accept a screening test if too many healthy horses are scratched from races. Strategies such as profiling and serial assessment may be required to appropriately identify a subpopulation at greater risk and thereby improve the positive predictive value of any applied diagnostic test.^10^

In this report, we evaluated use of DR and sCT imaging for injury prevention screening for condylar stress fracture. The limbs studied had MC3 PSG and condylar SBI with a range of severity from absent to severe that reasonably reflects Thoroughbred racehorse populations racing and training across the world. With DR imaging, there were often significant differences in assessment between observer and reference assessments of the extent of the dense subchondral bone, suggesting that it is more difficult for different observers to consistently identify mild to moderate adaptive change. Some observers had low sensitivity for detection of PSG subchondral fissure/lucency, suggesting assessment of the severity of the PSG lucency/fissure is also difficult with DR imaging. A similar trend was also found with sCT imaging but to a lesser degree. These conclusions were also supported by the overall low diagnostic sensitivity for detection of PSG subchondral sclerosis and PSG lucency/fissure with DR, which was improved with sCT.

Condylar stress fracture risk has been linked to subchondral plate functional adaptation by bone modeling,^26,40^ to increase the bone volume fraction and the thickness of the subchondral plate to better resist the high cyclic loads associated with racing. We found some bias in measurement of this variable amongst observers, suggesting that small differences in subchondral plate thickness are challenging to interpret clinically, particularly between observers. This might have been partially caused by measuring the thickness at slightly different reconstructed planes between observers. Like subchondral plate thickness, mechanical function has been linked to the presence of PSG subchondral lucency and its area in the sagittal plane.^30,31^ With sCT, there was good agreement between the observers and the reference assessment regarding whether there was a subchondral PSG lucency to measure. However, observer estimation of sagittal area was variable, and bias was identified between observer and reference measurements. This variation suggests that the usefulness of lucency sagittal area for risk assessment screening^30^ may be limited and that a more objective sCT-based risk assessment is needed.

Visual evaluation of the MC3 distal joint surface from 8 horses after cartilage digestion led to change in the risk assessment because a small amount of macroscopic fatigue damage was observed in the postmortem joint surface photographs but was not detectable because of the resolution of the clinical sCT. Findings were similar between horses. Most were also assessed as having normal risk by the panel of observers and it is acknowledged that the mildness of the changes may mean that their risk was low. The finding that not all horses with stress cracks arising from SBI in this study were detectable using either sCT or DR also suggests that serial assessment could be an important consideration in racehorse screening to detected SBI sooner rather than later before a horse with low risk of serious injury develops a progressive SBI associated with much greater risk. The experience of the authors suggests that SBI lesions and their associated risk can change rapidly over time.

Contralateral PSG subchondral lucencies and associated fatigue injury are known to be associated with catastrophically injured horses.^26^ The Thoroughbred used for assessment of intra-observer variation had contralateral condylar stress fracture at the time of euthanasia and the study limb was assessed as high risk with an overt stress fracture from fatigue injury that merited clinical intervention from the sCT evaluation that was then affirmed by the joint surface evaluation. We found that PSG lucency/fissure in this horse was inconsistently identified using DR amongst the observers and more consistently identified using sCT. One observer did not identify lucency/fissure in any of the repeats. For risk assessment, none of the observers rated any of the repeat image sets from this horse with the highest level of risk. Experimental mechanical testing data that includes this bone connects structural changes in subchondral fetlock bone to mechanical compromise,^30,31^ increasing confidence in the gold standard risk assessments in this study. To better inform clinicians regarding risk assessment for condylar stress fracture, the image sets used in this study have been made available via the Dryad data repository (datadryad.org) as a training set for veterinarians interested in fetlock risk assessment as part of injury prevention in racing Thoroughbreds.

For agreement with the reference assessment of MC3 subchondral structural changes, we found that the reliability of DR was generally poor (*r*<0.5). For sCT reliability was generally moderate (*r*≥0.50 to <0.75). Similar reliabilities were found when observer assessments were compared with each other. Whilst sCT assessments were generally more reliable, observer training may be important to ensure that any screening assessments are as accurate as possible.

Observers were generally better at detecting and assessing structural changes than risk assessment for imminent injury. Diagnostic sensitivity was lower than specificity for all observers and both imaging methods. We found diagnostic sensitivity for risk assessment was improved for all three observers who assessed both types of imaging when assessments were performed from sCT imaging. For horses determined to have a high risk of imminent injury by the reference assessment, observers generally underestimated risk. Whilst advanced imaging, such as sCT, has the potential to improve identification of at-risk horses, more data connecting structural change with mechanical compromise of the joint surface are needed to inform interpretation of diagnostic images.^31,41^

There were several limitations to this study. Assessment of fetlock diagnostic imaging is subjective and variable. Our study design has estimated this variation to better understand clinical usefulness of DR and sCT for fetlock screening. DICOM file sets were anonymized using the Horos DICOM viewer. One observer used Osirix for image viewing, which inadvertently grouped the repeated blinded image sets together, reducing blinding. This problem was addressed by reading the image sets as independently as possible. The library of samples we used appropriately represented the range of MC3 subchondral structural changes found clinically. However, the image set may not cover all potential clinical features or range of lesion severities. In this regard, intra-observer repeatability was only performed with one horse with PSG SBI to avoid making the data set used for analysis very large and reducing observer compliance. Intra-observer repeatability with different levels of subchondral bone structural change may be different. The presence of proximal sesamoid bone fracture in three limbs made evaluation of the MC3 subchondral bone more difficult, particularly one horse with biaxial comminuted mid-body displaced fractures. Such specimens were not excluded because the distal MC3 contained important subchondral pathology relevant to the study’s aims. Two of the four observers did not assess DR images from some of the horses because of concerns about image quality regarding resolution of the trabecular patterns in the MC3 distal metaphysis, and it was considered that the diagnostic quality of the DR images was of a generally lower level than that of best-practice *in vivo* imaging. Positioning of isolated limbs for flexed DP imaging is challenging relative to *in vivo* radiography. Furthermore, storage at −20°C and associated changes in the soft tissue compliance may also have affected image quality in this study. To address this problem, estimation of diagnostic sensitivity and specificity was limited to the horses that were assessed by all four observers, reducing sample size. In contrast, *in vivo* sCT imaging may be associated with more motion artifact with a consequent reduction in image quality, particularly if imaging is performed with the limb in a non-weight bearing position. Depending on the type of sCT system used clinically, imaging may be performed in a natural weight-bearing standing stance^25^ or with the limb suspended by ropes.^42^ Extension of this research approach to an *in vivo* image set is necessary to further assess DR image sets using optimal radiographic technique and sCT image sets obtained in the live horse. At UW-Madison, clinical experience with the Asto CT Equina^®^ for fetlock imaging suggests that there is little difference between *ex vivo* and *in vivo* imaging with this system as sCT images are made with the horse standing square with a natural stance. Finally, reference assessment of subchondral sclerosis was limited to analysis of sCT image data. These assessments could have been strengthened by objective assessment of the sclerotic volume in the MC3/MT3 bone end. Development of such a method is a current research priority.

In conclusion, we describe diagnostic sensitivity and specificity for detection of MC3 subchondral fatigue injury in the fetlock in an *ex vivo* model. Development of PSG SBI is a consequence of the high cyclic loads inherent to current Thoroughbred training and racing practices. Subchondral MC3/MT3 fatigue microdamage accumulates with age and cumulative race starts^43^ raising the possibility of targeted screening of horses in high-risk subgroups. Identifying when microdamage has reached a critical level requires more research and specific criteria for interpretation of subtle diagnostic imaging changes.^6,10,26,30,40^ Risk assessment through screening with diagnostic imaging is a promising approach to improve injury prevention in racing Thoroughbreds. However, the potential discrepancy between the presence of a subchondral stress crack identifiable pathologically and the typical diagnosis by diagnostic imaging through identification of a cortical discontinuity, fissure, or overt fracture line needs further study when diagnostic imaging is used for risk assessment as opposed to clinical diagnosis. Use of sCT imaging for screening of individual horses is currently being used by Racing Victoria for the Spring Racing Carnival (www.racingvictora.com.au). Knowledge of sensitivity and specificity of fetlock lesion detection by DR and sCT is critical information needed to develop improved screening programs for racehorses and to begin to establish appropriate roles for DR and sCT imaging in injury prevention in Thoroughbreds. Our results show improved detection of MC3 subchondral structural change and risk assessment for condylar stress fracture with sCT in an *ex vivo* model and contribute to improved understanding about how diagnostic imaging may best be used to detect SBI that may predispose to serious or catastrophic condylar stress fracture.

## SOURCES OF FUNDING

This study was funded in part by a grant from the Grayson Jockey Club Equine Research Foundation.

## SUPPLEMENTARY FILES

**Supplementary File S1**. Fetlock imaging assessment form template.

**Supplementary File S2**. Supplementary figures and tables.

**Supplementary File S3**. Supplementary data.

## Author contributions

S. Irandoust contributed to study design, data collection, data analysis, interpretation, and preparation of the draft. L. O’Neil contributed to data analysis and manuscript figure preparation.

C.M. Stevenson contributed to data collection. F.M. Franseen contributed to data collection.

P.H.L. Ramzan contributed to the study design, data collection, and interpretation of the results.

S.E. Powell, S.H. Brounts, and S.J. Loeber contributed to data collection, and interpretation of the results. D.L. Ergun generated the sCT images for the project. R.C. Whitton contributed to the study concept and manuscript writing. C.R Henak contributed to study design, data analysis, and interpretation of the results. P. Muir conceived the study, contributed to study design, data collection, data analysis, interpretation, and preparation of the draft manuscript. All authors approved the final version of the manuscript.

## Conflict of interest

P. Muir is a Founder of Asto CT, a subsidiary of Centaur Health Holdings Inc. and a founder of Eclipse Consulting LLC. S.H. Brounts is a clinical advisor to Asto CT. D.L. Ergun is the Chief Executive Officer of Asto CT.

## Ethical animal research

All data is from analysis of *ex vivo* limbs.

## Acknowledgements

We gratefully acknowledge the help of all the Comparative Orthopaedic Research Laboratory students who contributed to this project. We are grateful to Brett Nemke and Henry Benchimol for helping with collection of radiographs. We also are grateful to Dr. Patricia Marquis who contributed limb specimens for analysis.

## Data accessibility statement

The data that support the findings of this study are available in an online data set from the Dryad Data Repository at https://doi.org/10.5061/dryad.dncjsxm55.

## Notes

https://doi.org/10.5061/dryad.dncjsxm55

