## Supplementary File S2 for "Comparison of radiography and computed tomography for condylar fracture risk assessment in Thoroughbred racehorses"

**Comparison of planar digital radiography and helical standing computed tomography for assessment of condylar stress fracture risk in Thoroughbred racehorses**

S. Irandoust,^1,2^ L. O’Neil,^1^ C.M. Stevenson,^1^ F.M. Franseen,^1^ P.H.L. Ramzan,^3^ S.E. Powell,^4^ S.H. Brounts,^1^ S.J. Loeber,^1^ D.L. Ergun,^5^ R.C. Whitton,^6^ C.R. Henak,^2,7,8^ P. Muir^1,*^

^1^Department of Surgical Sciences, University of Wisconsin-Madison, Madison, WI, USA

^2^Department of Mechanical Engineering, University of Wisconsin-Madison, Madison, WI, USA

^3^Rossdales Veterinary Surgeons, Newmarket, UK

^4^VetCT, Cambridge, UK

^5^Asto CT, 7921 UW Health Ct, Middleton, WI 53562, USA

^6^Department of Veterinary Clinical Sciences, Faculty of Veterinary and Agricultural Sciences, University of Melbourne, Werribee, Victoria, Australia

^7^Department of Biomedical Engineering, University of Wisconsin-Madison, Madison, WI, USA

^8^Department of Orthopedics & Rehabilitation, University of Wisconsin-Madison, Madison, WI, USA

Author contributions:

S. Irandoust contributed to study design, data collection, data analysis, interpretation, and preparation of the draft. L. O’Neil, . C.M. Stevenson contributed to data collection. F.M. Franseen contributed to data collection. P.H.L. Ramzan contributed to the study design, data collection, and interpretation of the results. S.E. Powell, S.H. Brounts, and S.J. Loeber contributed to data collection, and interpretation of the results. D.L. Ergun generated the sCT images for the project. R.C. Whitton contributed to the study concept and manuscript writing. C.R Henak contributed to study design, data analysis, and interpretation of the results. P. Muir conceived the study, contributed to study design, data collection, data analysis, interpretation, and preparation of the draft manuscript. All authors approved the final version of the manuscript.

Conflict of interest:

P. Muir is a Founder of Asto CT, a subsidiary of Centaur Health Holdings Inc. and a founder of Eclipse Consulting LLC. S.H. Brounts is a clinical advisor to Asto CT. D.L. Ergun is the Chief Executive Officer of Asto CT.

Ethical animal research:

All data is from analysis of *ex vivo* limbs.

Acknowledgements:

We gratefully acknowledge the help of all the Comparative Orthopaedic Research Laboratory students who contributed to this project. We also are grateful to Dr. Patricia Marquis who contributed limb specimens for analysis. The authors are grateful to Professor Chris Whitton, University of Melbourne for advice regarding study design.

Data accessibility statement:

The data that support the findings of this study are available in an online data set from the Dryad Data Repository at https://doi.org/10.5061/dryad.dncjsxm55.

**Supplementary Figures and Tables**

**Supplementary Figure S1**. Distribution of joint pathology scores from assessment of articular cartilage and subchondral bone from the distal end of MC3 in Thoroughbred racehorses. Damage to the cartilage and subchondral bone ranged from absent to severe in the sample population. LC – lateral condyle, LPSG – lateral parasagittal groove, MPSG – medial parasagittal groove, MC – medial condyle.

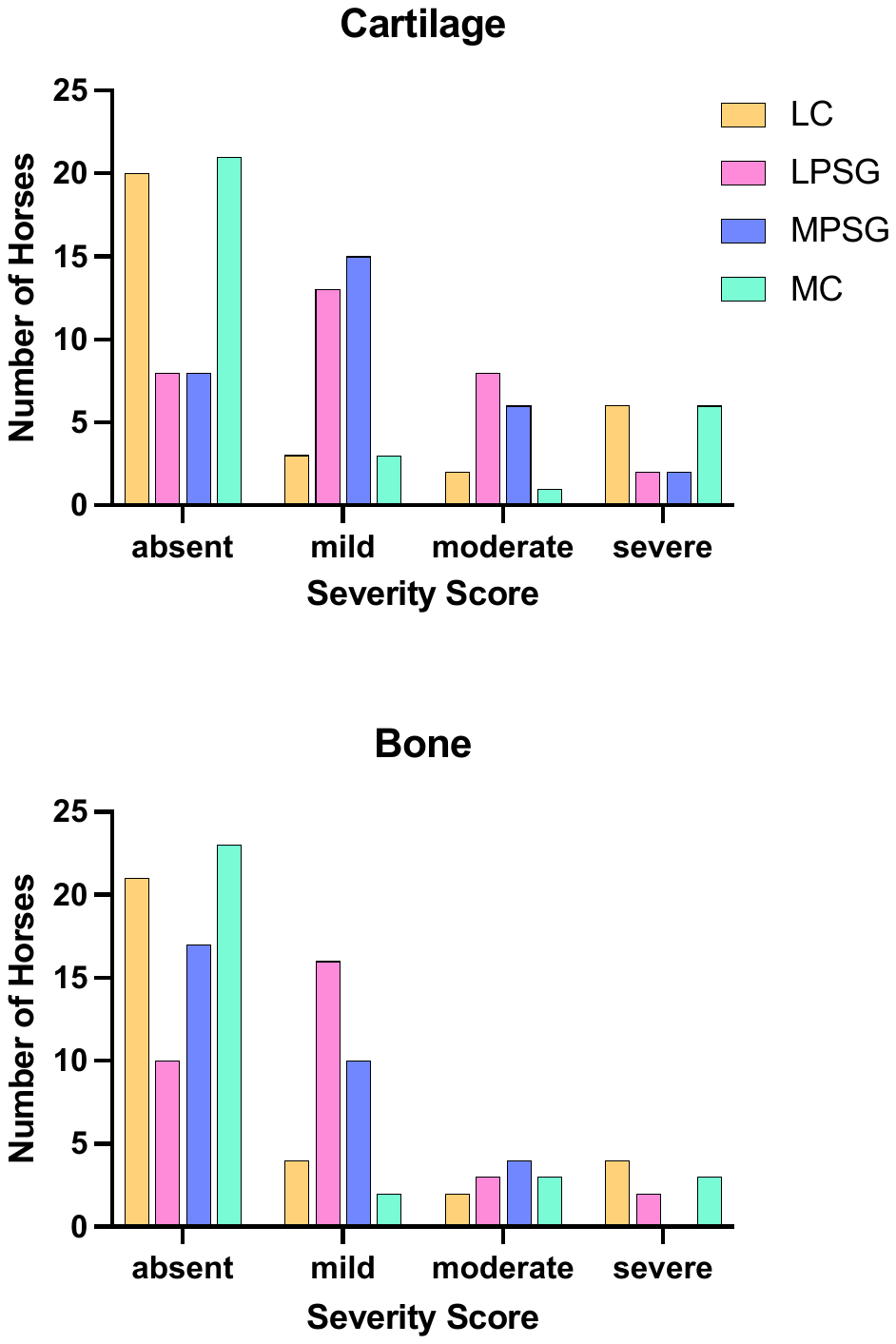

**Supplementary Figure S2**. Image montage presenting the surface appearance of the subchondral bone plate of the distal end of each third metacarpal bone after digestion of the articular cartilage for the 31 racing Thoroughbreds studied. **Note**. Lateral is to the left and palmar to the bottom.

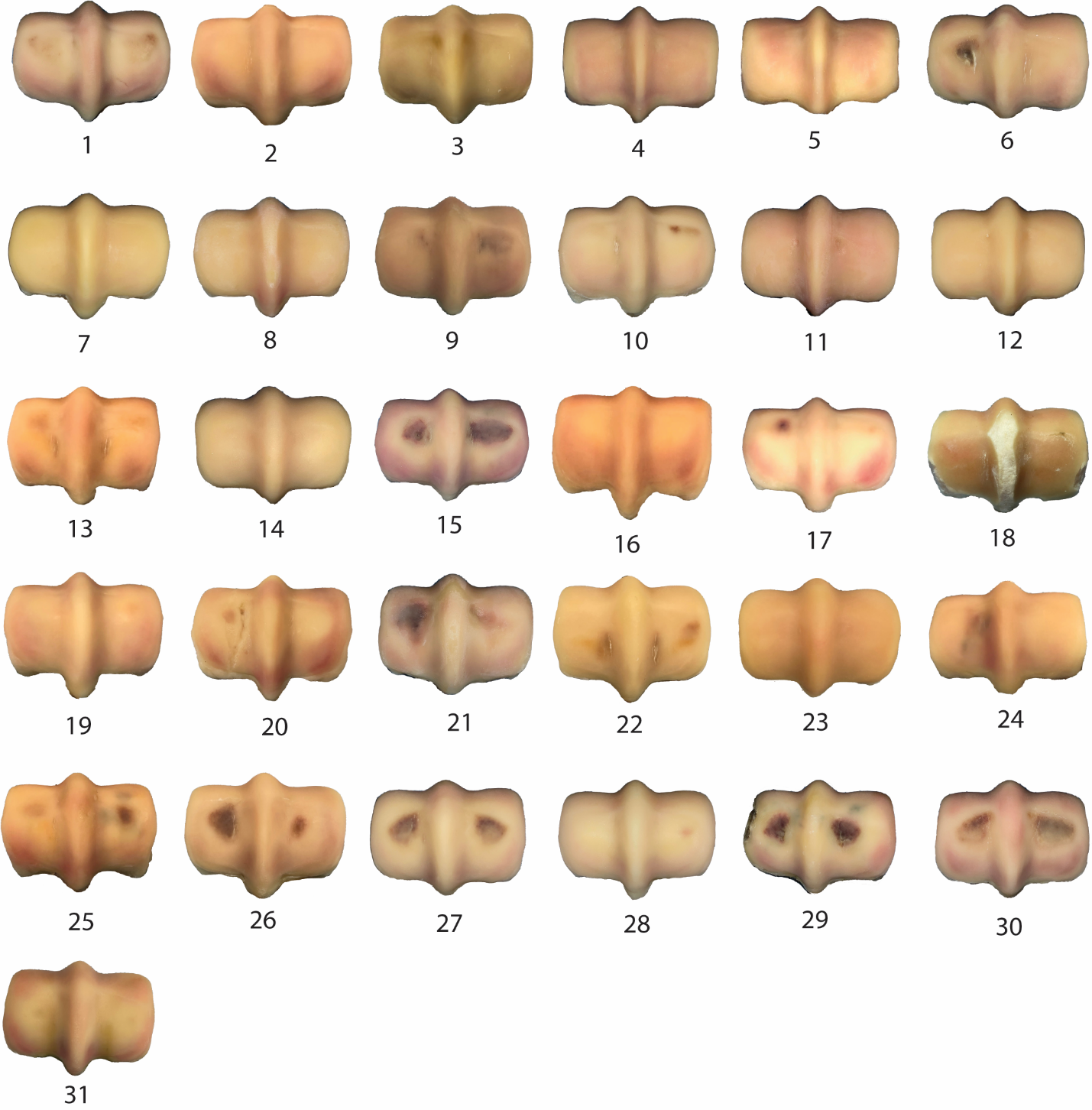

**Supplementary Figure S3**. Bland-Altman plots for measurement of parasagittal groove subchondral plate thickness from standing computed tomography imaging in Thoroughbred racehorses. Measurements by the observers showed positive (Observer 1) or negative bias (Observer 3) from the reference measurements.

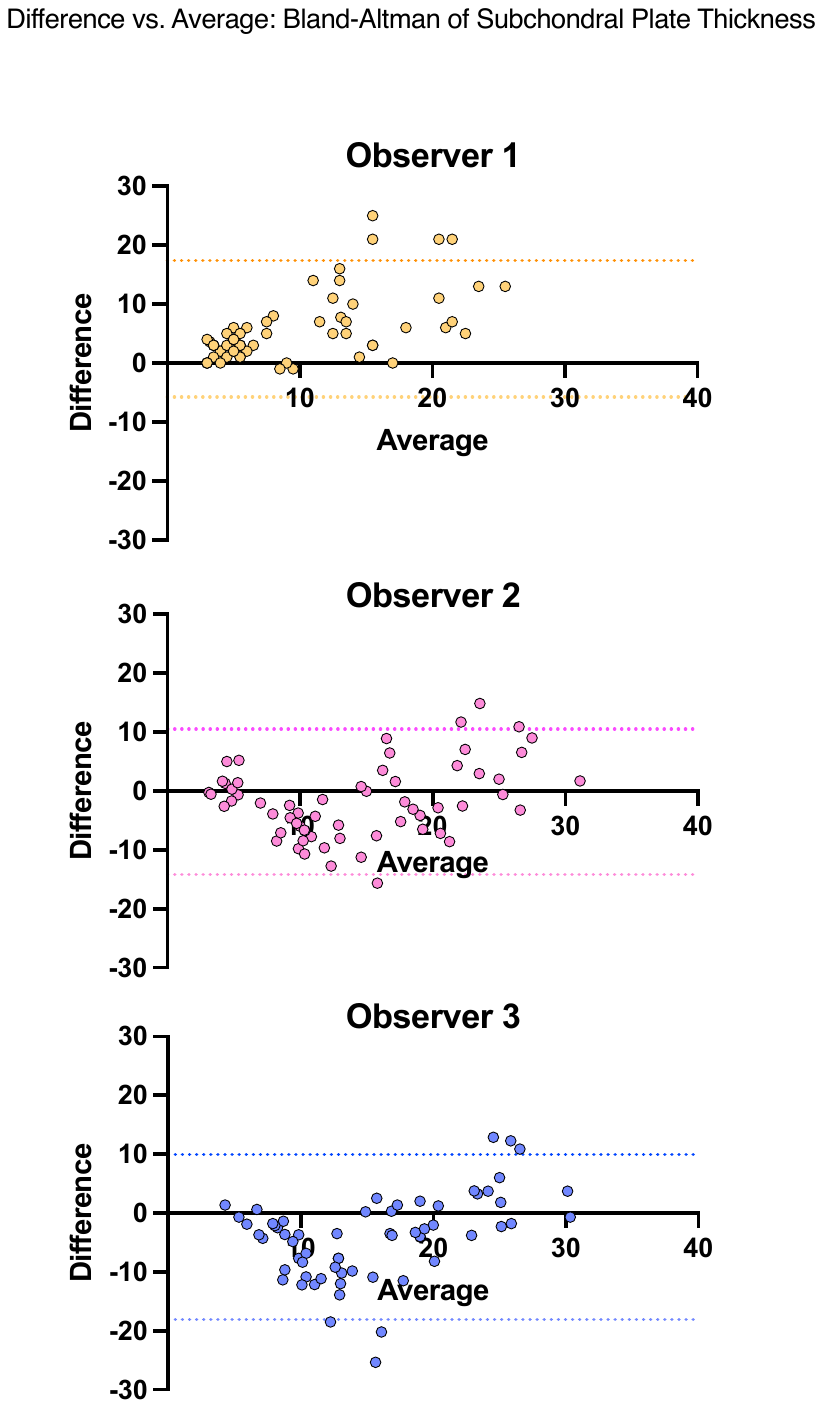

**Supplementary Figure S4**. Bland-Altman plots for measurement of parasagittal groove subchondral lucency area (mm^2^) in the sagittal plane from standing computed tomography imaging in Thoroughbred racehorses. Measurements by Observers 1 and 3 showed positive bias from the reference measurements. The data points included in the plot were those with measurements from both the observer and the reference assessment. This resulted in a sample size of 14, 16, and 9 for Observers 1 to 3, respectively.

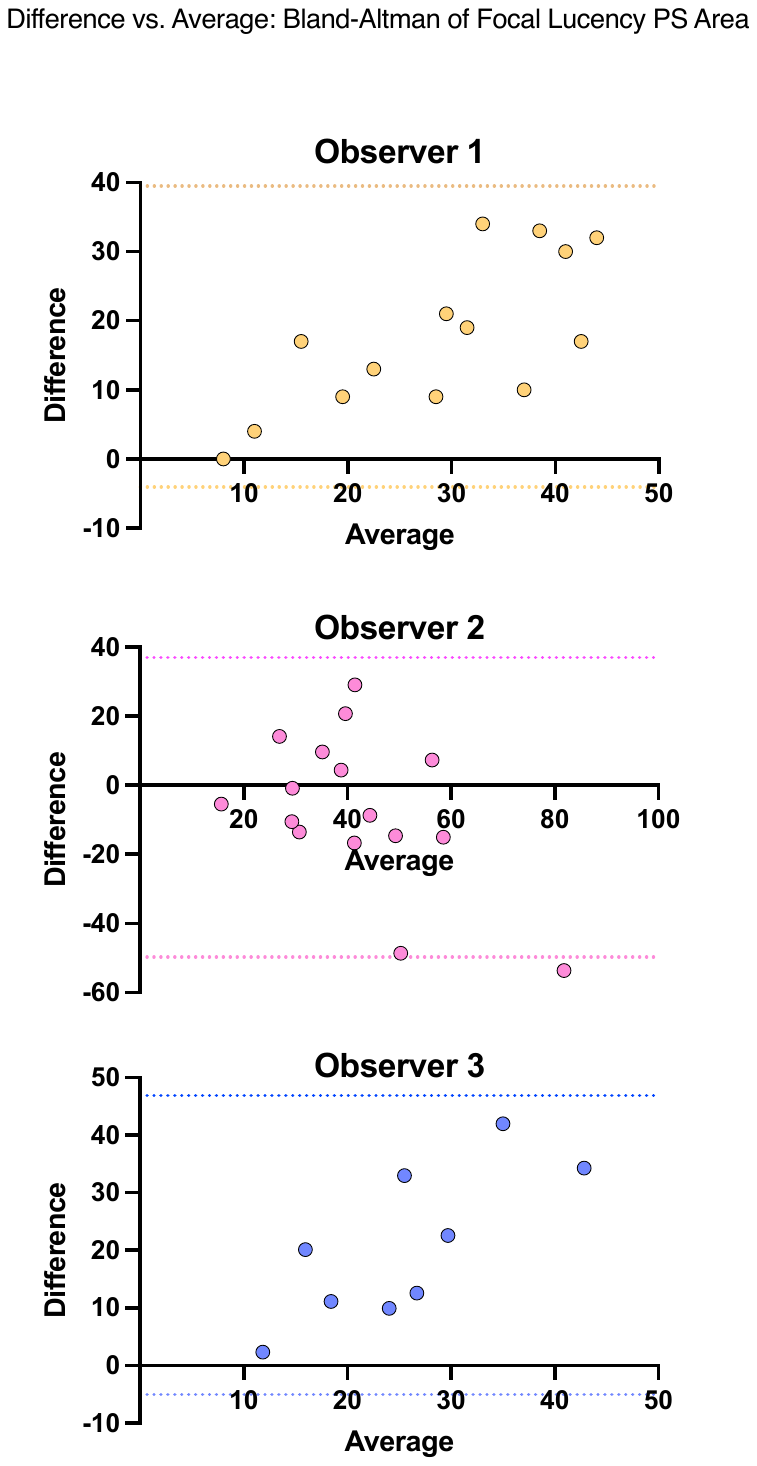

**Supplementary Figure S5**. Postmortem (**A**), digital radiography (**B**), and standing computing tomography (sCT) images (**C-G**) from a racing Thoroughbred with a lateral parasagittal groove (PSG) focal subchondral bone lucency because of fatigue injury (Horse #10, Figure 1). The lucency seen on sagittal (C), transverse (D), and dorsal (E) sCT reconstructions correspond to the linear PSG subchondral fatigue cracks seen in A. The reference assessment was *Overt stress fracture from fatigue injury.* A lower risk assessment rating was provided by all 4 observers interpreting DR imaging. Concordance with the reference assessment was improved with sCT imaging. **Note**. Lateral or dorsal to the left.

**TABLE S1**. Observer variation in detection of sagittal plane parasagittal groove subchondral bone lucency in Thoroughbred racehorses using standing computed tomography

|  |  | **Reference Assessment** | |
| --- | --- | --- | --- |
|  |  | Yes | No |
| **Observer 1** | Yes | 14 | 2 |
|  | No | 5 | 40 |
| **Observer 2** | Yes | 16 | 4 |
|  | No | 3 | 36 |
| **Observer 3** | Yes | 9 | 5 |
|  | No | 2 | 30 |

**Note**. The sample sizes for each observer varied because some observers made measurements for lesions that were considered to occupy both the condyle and parasagittal groove regions. For consistency, results are reported for PSG lucencies only. The number of measures for each observer was: Observer 1 - 61, Observer 2 - 59, Observer 3 – 46.
